# Design and Implementation of a Decision Making System for Controlling a Hand Exoskeleton Based on EEG/EMG Signals

**DOI:** 10.1101/2025.09.10.675353

**Authors:** Alireza Zolanvari, Maziar Arfaee, Mohammad Bagher Menhaj, Vito De Feo, Nabiolah Abolfathi, Fatemeh Jahangiri, Ahmad Sohrabi, Atena Sajedin

## Abstract

This paper presents an approach of combining Electroencephalography (EEG) and Electromyography (EMG) signals to create a hybrid Brain Interface Computer (BCI) device for controlling a hand exoskeleton through classifying the flexion attempt and resting states of the subjects. We analyzed data of 51 healthy and patients with brain lesions that involved the motor cortex and different hand movement disorders. Their EMG and EEG activity were recorded while the subjects attempted to move their index finger. The signals are analyzed through deep neural network, restricted Boltzmann machine and LDA classifier methods to access a reliable accuracy. Further, to evaluate the proposed approach, we designed and implemented a robotic system for rehabilitation of the hand movement to show that it is able to derive assistive control, by detecting flexion movement from the signals as a command and send it to robot. The system is combined with control signals that determine the information of flexion according to measured EMG signals.

## Introduction

Exoskeletons are great wearable mobile systems supporting mechanical benefits to the users. People with neuromuscular disorders like spinal cord injury, amyotrophic lateral, sclerosis brainstem stroke, cerebral palsy or multiple sclerosis, encounter a severe decline in quality of life and a significant burden on family and society. Therefore, the efforts should be assisted them with different types of mobility and accurate communication systems [Wolpaw etal,. 2002]. However, since these systems must interact directly with disabled human users, their design and performance requires more sophisticated methods and technologies which have higher accuracy, reliability and safety. The Brain Computer Interface (BCI) system has found interesting applications in the field of bio-robotics. BCI allows patients to directly use their brain activity (electroencephalography (EEG) signals) to control automated systems, especially in several bio-applications [Al-Quraishi et al., 2018; H. Gao et al., 2019; Kucukyildiz et al., 2017; Lee et al., 2017; Li et al. 2018 and Murphy et al., 2017, Noormohamadi et al.,2014, Mirzabagherian et al., 2023]. EEG and local filed potentials reflect the behavior of cortical neurons and the components of these signals different bands (e.g. delta, theta, alpha, beta or gamma bands) are provide a physiological correlate of cognitive phenomena [Pesaran et al., 2002; Gail et al., 2004; Fries et al., 2008; Sajedin et al., 2019; Sajedin et al.,2024, Mitzdorf, 1985] and have been suggested to play an important role in cortical processing.

High temporal resolution and portability of EEG based data acquisition makes it a popular choice in developing neuro-rehabilitation applications [Prakash, et al., 2021]. However, EEG-based systems are not fully suitable for bio-robotics applications as their low reliability and low user compatibility [Wolpaw et al., 2002; Wolpaw, et al., 2000; Chowdhury, et al. 2017]. As a result, to overcome these problems, taking advantage of both Electromyography (EMG) and EEG-based control strategies has become a promising technique to structure a hybrid system that making the overall system more accurate and reliable [Leeb et al., 2011, 2010; and Kiguchi and Y. Hayashi, 2018]. Several findings illustrated that using EEG and EMG signals simultaneously improve the performance of the system performance [Leeb et al., 2011; Dulantha Lalitharatne et al. 2013; Wöhrle et al. 2017; Tryon et al. 2019; Tortora et al. 2020; Tryon and Trejos 2021]. This approach improves classification accuracy as well as reliability, by simultaneously leveraging the benefits of the signals. Studies have shown that EEG–EMG models reach to acceptable accuracy even during EMG signal attenuation influenced by muscle fatigue [Leeb et al. 2011; Tortora et al. 2020; Li et al. 2017], illustrating the increased reliability that obtained through using multiple signal types simultaneously. Many findings were illustrated that the accuracy of the system increases by using hybrid approach as compared to used EMG and EEG signals separately [Leeb, et al., 2010 and 2011; Kiguchi et al. 2012]. Moreover, in another studies robotic arm is controlled by hybrid BCI system [Kiguchi et al. 2014; Salgado et al.2016].

Numerous studies have reported and reviewed the effectiveness and even necessity of exoskeletons and hybrid rehabilitation robots [Borboni et al., 2017; Dovat et al., 2010]. The important reason is the possibility of performing repetitive and precise movements as well reducing the duration and cost of treatment [Huang et al. 2018; Iqbal et al. 2014]. Initially, the lower limbs, due to the size and simplicity of their joints, received more attention from scientists [Fu, et al. 2008], and today these exoskeletons have reached a significant level of development. But, due to the complex structure of the hand and its high capabilities in doing things, implement of hand exoskeletons has become very challenging [Borboni et al., 2017; Rahman and Jumaily, 2012]. In recent years, many hand rehabilitation systems have been developed, to helps hand movement of the people with spinal cord injury [Ueki et al. 2010] or a virtual reality-enhanced rehabilitation system for stroke patients [Vanoglio et al., 2018]. Different upper-limb rehabilitation devices, including exoskeleton gloves are reviewed in [Maciejasz et al., 2014]. However, these rehabilitation devices do not guarantee complete recovery of the motor-sensory functions of the affected limb. Up to now, several studies have been addressed different aspects of exoskeletons, such as architecture, system design, prediction, and control system. However, most works have focused on a specific element of design or application without providing a comprehensive framework. Therefore, we aimed to present a comprehensive study, and propose an architecture to provide motion active assistance, based on EEG and EMG signals, and design and implement a rehabilitation system.

In the present study, we analyzed data of 51 healthy subjects and patients with brain lesions that involved the motor cortex and different hand movement disorders. Their EMG and EEG activities were recorded while the patients attempted to move their index finger. Different approaches were tested to detect the movement intentions of the subjects including restricted Boltzmann machine, Deep belief neural network, Sparse auto encoder network and (linear discriminative analysis) LDA classifiers. Then, as a hardware interface, we designed and implemented a glove-type exoskeleton, with a soft structure instead of rigid links and joints which allows the device to be compact. The proposed hand exoskeleton is light, comfortable, and portable, and both hardware and training paradigm were considered during the design process to receive and execute the flexion movement commands from EEG and EMG signals. For this purpose, we developed a control architecture to provide motion active assistance through EMG signals that providing complementary information to control commands, and as soon as the quality of these signals decreased, the system switch to communicate the results of EEG signals processing to the hand robot and the actuators. The objective of this work is to present a method of combining EEG and EMG signals to create an accurate hybrid BCI device for controlling a hand exoskeleton through classifying the flexion attempt and resting states of the subjects. This work provides an example of EEG–EMG fusion through different classifier, and highlights the most promising methods to use for further development.

## Experimental Procedures

### Subjects

Fourteen healthy subjects and thirty seven patients were considered in this study. The patients had suffered different disorders like Emiplegics, Aphasics and Ataxics that involved the motor cortex. The subjects sat in a chair looking at a LCD monitor placed approximately one meter in front of them. Subjects were instructed to observe a timer and perform a brisk finger (index) flexion every 10s starting from 5s, 40 finger flexions were performed for a session of 400.

### EEG/EMG Recordings

EEG was recorded by 7 monopolar scalp Ag/Ag/AgCl electrodes according to the standard 10-20 system referenced to the ground over A1 and A2. The EEG was amplified, bandpass filtered between 0.015 Hz and 50 Hz, and then sampled at 512 Hz with a time constant of 0.1 s. Surface EMG was recorded from the right hand palm by means of two electrodes, one above the then areminence and the other one in correspondence to the first joint of the index finger. sEMG was amplified, bandpass filtered between 5 Hz and 500 Hz and then sampled at 512 Hz with a time constant of 0.3 s. Both EEGs and sEMGs were notch filtered at 50 Hz to remove the power line noise. The impedance of each electrode was kept above 5 Kohm. EMG, data presents 2channels recording that refers to the electrode in the thenar eminence and the first joint of the index finger. For the experiment we used EMG1-RF that recorded over the index.

### Signal Processing and Data Analysis

Due to the limitation of the portability of the robot, and the limitation of the processing power of the central processor, we extracted the appropriate characteristics of the signals with the minimum amount of calculations. Different types of features were extracted by Root Mean Square (RMS), Waveform Length (WL), and Log detector (LOG) from the EMG signals. For EEG signals, two methods of feature extraction of wavelet and FFT were used. Feature extraction was performed in windows of 250 ms, with sliding window of 125 ms. Since the brain signals are non-stationary, we utilized the Restricted Boltzmann Machine based method, which has the ability of learning a probability distribution. In addition, since deep neural networks can efficiently and completely modeled the most common functions, especially learning functions, we also used deep neural network by cascading several RBM networks named Deep Belief Network (DBN).

### Restricted Boltzmann Machine

Restricted Boltzmann machines (RBMs) are a generative probabilistic artificial neural network with the ability of learning a probability distribution. RBMs are applied for dimension reduction, feature extraction and modelling. RBMs are special class of Boltzmann machines, with the restriction that their neurons should form a bipartite graph in which a pair of nodes from each two groups of units may have a symmetric connection between them. Also, there are no connections between nodes within a group. This restriction leads to more efficient training algorithms than existed for the general form of Boltzmann machines. Computational power enhancement and the development of faster learning algorithms made RBMs suitable to the machine learning problems. The other advantage of the RBM method is that it can hold the information of the input vector and their label in the hidden units, and therefore, without the need to determine the threshold, the input is classified by the network. RBMs proposed as building blocks of multi-layer learning systems called deep belief net-works. The RBM structure used in this study contains 21 observable layers, and 10 hidden layers. The other parameters are given in Table 1.

**Table 1.** RBM Parameters.

| Parameters | Values |
| --- | --- |
| Layers number | 11 |
| Nodes of hidden layers | 10 |
| Batch size | 100 |
| Learning rate | $10^{-4}$ |
| Maximum number of iterations | 50 |

### Deep Belief Network

By cascading several RBMs network, a network was made which named Deep belief network (DBN). The main feature of this network is the ability of processing the input layer by layer in order to reach a better predictive model in each step. So, after finding the best connection weights for the RBM in the first layer, we can use it as the initial value of the weights in the second layer and be sure to have a better result than the previous step (see Figure 1). Where, the i^th^ layer training data hi, is the data that related to the features extracted from the previous layer. This layer-by-layer method is repeated several times to obtain a deep hierarchical model. The input data v is the observable variables.

**Figure1.**
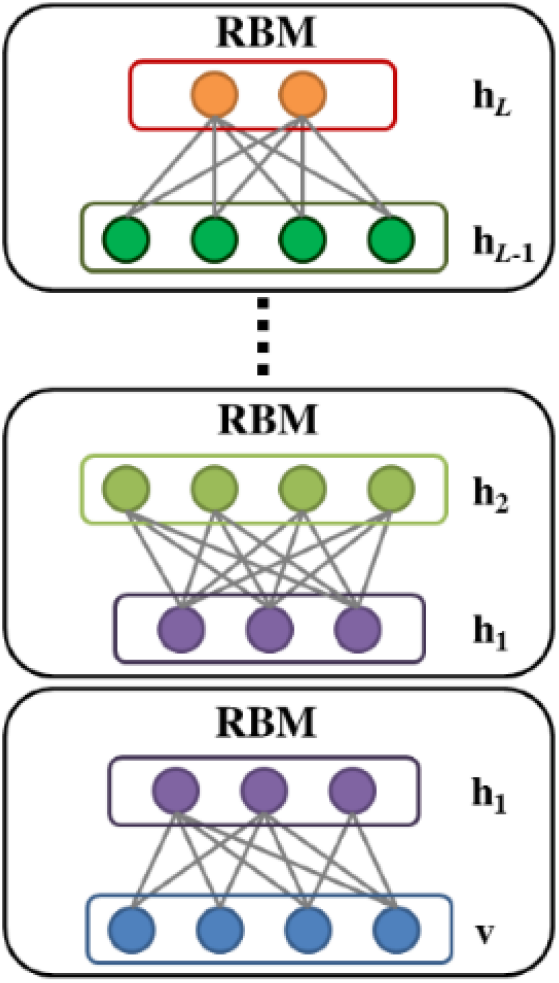
Scheme of a Generative Deep Belief Network (DBN)

DBN is a generative network which attempts to predict the value of its sets of input vector through the trained model. Thus, it is usually used for unsupervised learning procedures. We performed a binary classification task using this network based on the reconstruction error. Therefore, we trained the network using the samples of one of the classes, and tried to minimize the reconstruction error for the signals of that specific class. Then, using a hyperplane the signals with low reconstruction error are classified in one class and others are classified in the other one.

### Discriminative DBN

If we compose a DBN using multiple discriminative RBMs, the established network is called Discriminative DBN (Figure 2). Unlike general DBN, it is common to apply this discriminative network to supervised learning and classification tasks. The directionless activity of the DBN is divided into two parts: encoding and decoding as seen in Figure 3. This method is also called self-encryption, since the input signal is received by the network and then reconstructed. After that, by comparing the original signal and the reconstructed signal with the threshold, the signal is classified.

**Figure 2.**
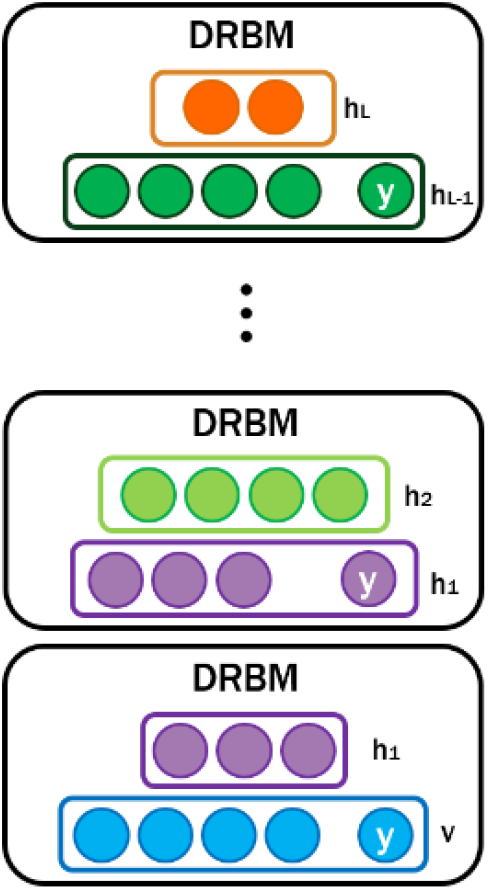
Scheme of a Discriminative Deep Belief Network

**Figure 3.**
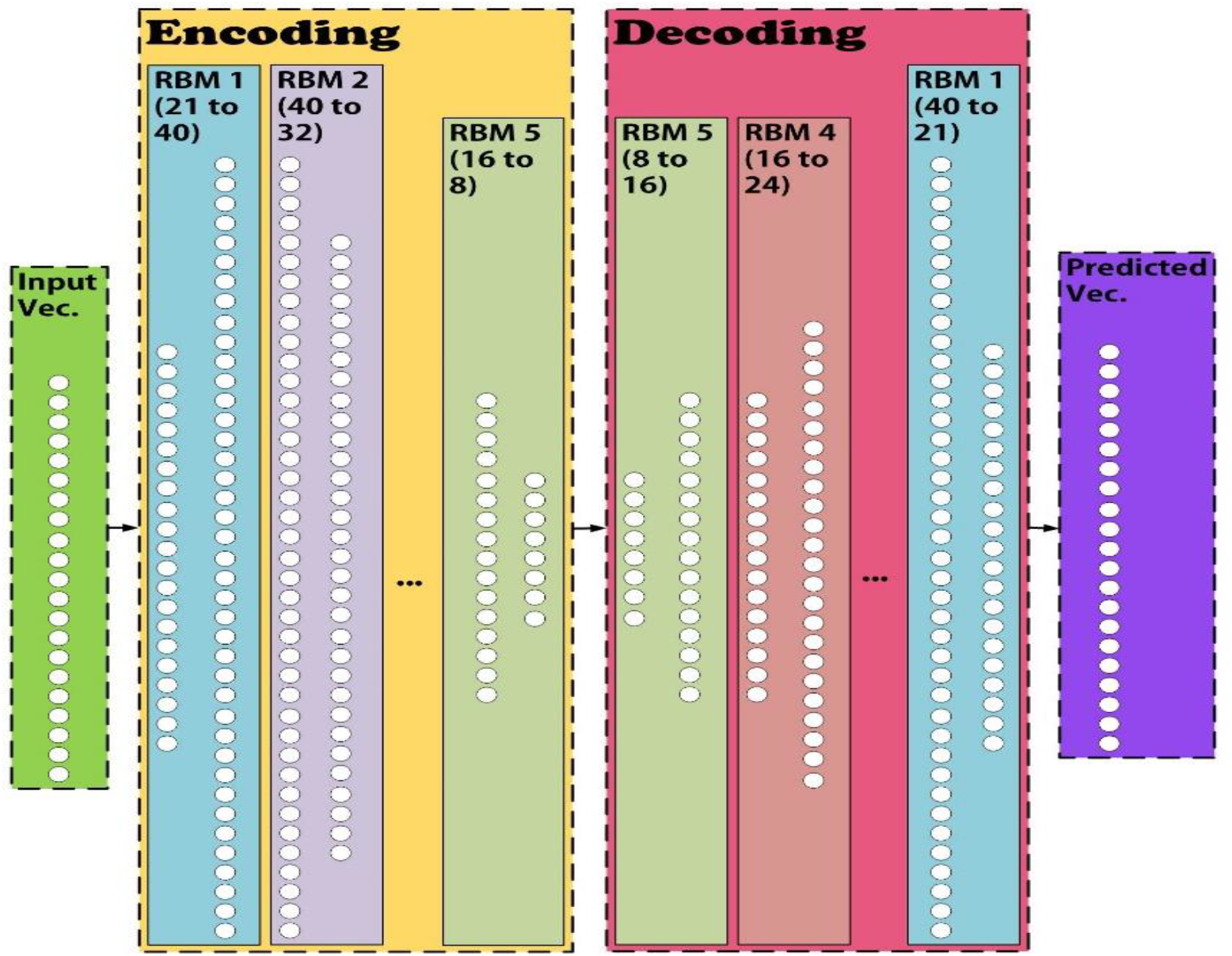
The Architecture of Deep Belief Network (DBN) The networks contain 11 hidden layers each included 40 to 8 neurons respectively.

We utilized DBN, for both feature extraction and model prediction of the bio-signals. In order to set the neural network, we determined the important parameters such as number of the layers, number of the neurons in each layer, the activation function of neurons, and the learning rate of the network. The value of these parameters was obtained by try and error method. As a result, Figure 3 showed the employed networks which contained eleven hidden layers, each included 40 to 8 neurons, respectively. In order to reduce the complexity of the network, we used the RelU activation function in the hidden layers. More details of the network architecture are given in the Table 2.

**Table 2.**
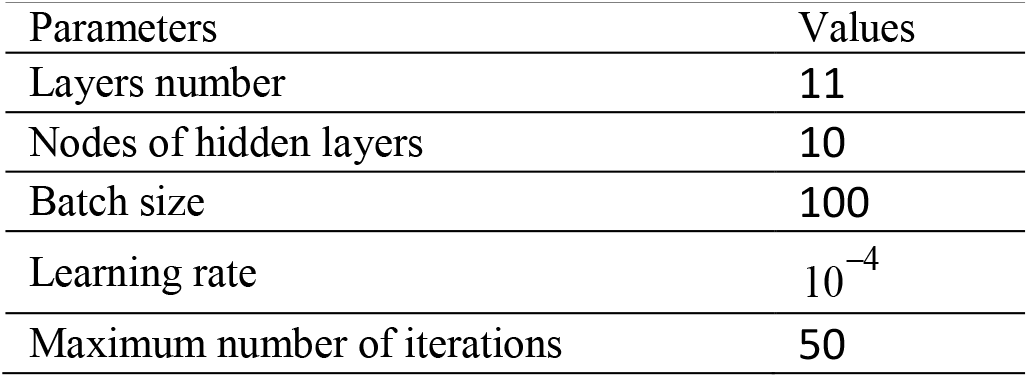
Deep Belief Network Parameters.

### Parallelizing Two Generative DBNs

Here, we described the hybrid schemes that we suggested based on DBNs to classify the bio-signals. The training data is recorded in a condition that the subject didn’t have any extra movement, but in the real life, the user may have many other activities while they want to move their fingers. Here, we addressed this problem by parallelizing two Generative Deep Belief Networks (Figure 4). In this method, each network tries to figure out the model of each class separately. Therefore, the input signal reconstructed by both networks, and the sample is related to the class which has less reconstruction error.

**Figure 4.**
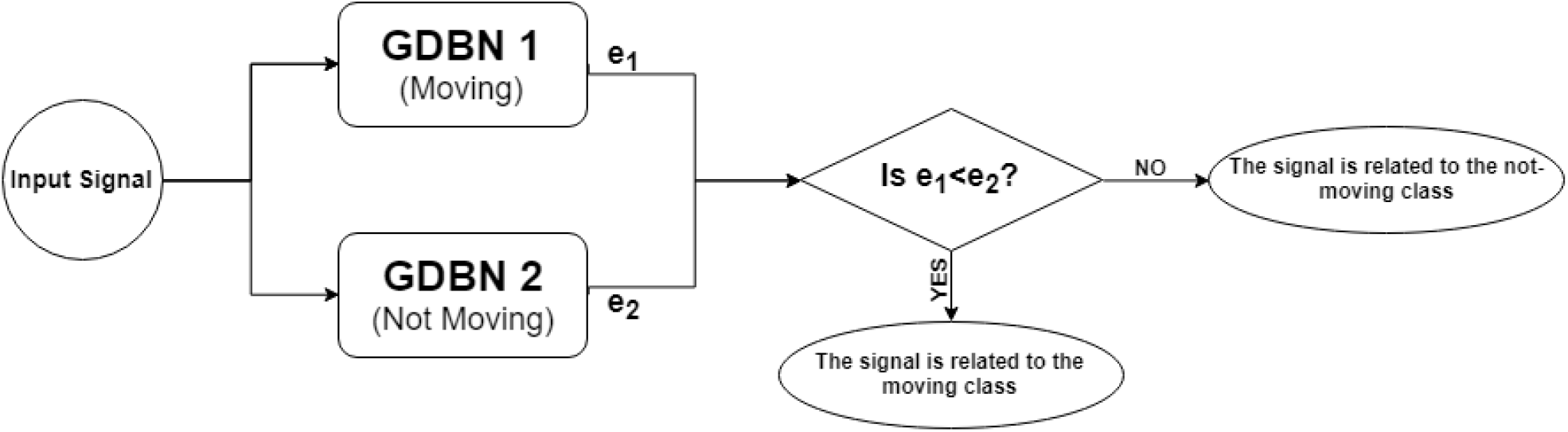
Block diagram of the classification using two parallel GDBN

### Cascading Two Parallelized GDBN to an LDA Classifier

At the previous section, allocating each sample to its related class has been done according to a hard threshold which had been chosen through try and error method. In order to choose this threshold more accurate, we utilized LDA classifier. This classifier found the discriminative bound automatically according to its inputs which are the reconstruction error of each GDBN. Where, p_1_ and p_2_ are the membership probability of the input respected to the first and the second class (Figure 5). Notably, in all the evaluations, 80% of the brain signals were used for training and the remaining 20% were used for the evaluation.

**Figure 5.**
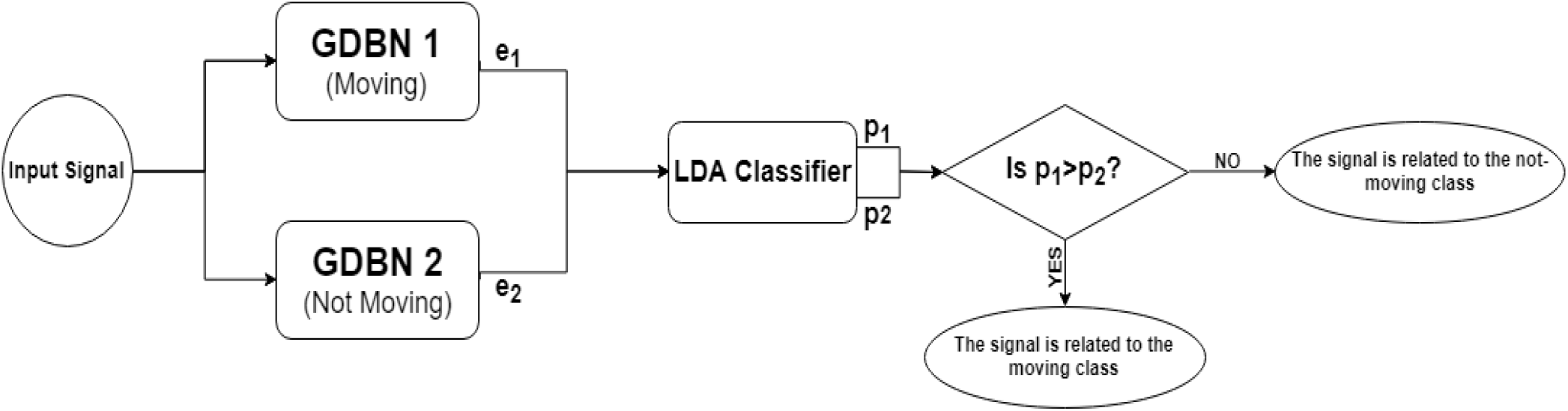
Block diagram of the classification afterascadingan LDA classifier

## Hardware interface setup

In order to validate the developed model, we designed and fabricated a wearable hand robot by which it is also possible to employ the processed data to move disabled fingers. The robot is driven by thin metal cables passed through the palm to the finger tips and phalanges. A major drawback of wearable robots with rigid links is high probability of misalignment between the joints of a robot and native joints of human. Using cables instead of rigid link to actuate the robot is beneficial as it is flexible and can solve the misalignment problem that is prevalent in robots with rigid links [Khomami, et al. 2019]. In this system, each finger is moved by two cables, each responsible for different movements. On each finger, one wire is connected to the fingertip, which moves the DIP and PIP. Second cable is connected to MCP and is responsible for flexion around MCP joint. Five servo-motors (Dynamixel XL-320) are employed to pull and release the cables, one for each finger. An adjusting system is designed for tuning the effective length of cable. Therefore, it enables us to have desirable movement.

The robot is active in flexion movement and passive in extension movement. Five motors are used for flexing the fingers in this robot, one motor for each finger. However, the extension of each finger is done using a tension spring. Finger extension movements are only for opening the hand and there is no need to apply any force in this movement. This reduces the number of actuators and mechanisms and therefore the weight of the robot. Flexion movement of each finger is done by a small and light-weight servo motor (Dynamixel XL-320) that has an integrated gearbox which makes it capable of providing enough force despite of its small size. The motors are placed on a plate that is fixed on top of the hand. Due to the small size and light weight of the motors, we are placed on a plate that is located on top of the hand.

For the thinness and strength of the cables, filament cables consisting of 49 thin metal strands with a diameter of 0.5 mm have been used. The cables pass through a spring sheath with an outer diameter of 1.8 mm and reach the fingers. The cables are connected to the finger joints by a very small piece. These parts are attached to the finger joints on the glove in such a way that the force is applied to a point that creates the most torque and the least vibration. The required power of the motors as well as the CPU must be supplied by the battery due to the portability of the robot. The battery used must be able to supply 7.4 volts nominally as well as a nominal current of 1.1 amps. In addition, due to other constraints such as the safety of the battery under pressure in certain conditions, lightness and no deformation over time, we used two series of cells made of lithium-ion battery. Size of the dual-cell battery with an electric charge of about 2600 mAh are about 35 x 35 x 100 cm3, which with the CPU, was attached to the user’s arm in the form of an armband.

We chose the Orange Pi One single-board computer because of its small size, light weight, high processing power, and cost-effectiveness. For the simplicity and lightness of the robot, to connect the robot to the hand, we sew it to a glove. Due to their elasticity, gloves can adapt to a limited range of different hand sizes.

## Results

### Filter and Inputs

Motor imagery preparation for movement or actual movement is usually observed by decrease in low frequency rhythms over the sensorimotor cortex area, especially the contra-lateral region. The statistical characteristics of the rhythms such as power, variance and logarithm were extracted by deep neural network. For each EEG channel, three filtered EEG bands were generated: 7-15 Hz, 15-25 Hz, and 25-30 Hz. We have seven EEG channels data. Therefore, our input for the deep network is an N× 21 matrix if we gathered N samples per each user.

In addition, during the recording brain signals, there are the possibility of unwanted movements such as eye movements, contraction of the jaw muscles and similar movements. This information can contaminate the signal and cause errors in the next processing steps. Therefore, by analyzing the signal, it is possible to obtain the amount of signal pollution and refine it. Simulations were run in Matlab sofware.

### Performance Evaluation

For classifying signals into two classes “Moving” and “Nonmoving”, we performed and compared four types of classifiers that are described in the Method section. The performances of these algorithms are compared using two main metrics that are Cohen’s Kappa and Accuracy.

Figure 6 illustrated a comparison between the performance of DBN, discriminative DBN (DDBN), two parallelized DBNs (PDBN), and the combination of parallelized DBNs and LDA (PDBN+LDA). In order to have a better insight for comparing these algorithms for classification task, the result of confusion matrices and ROC curves are presented in the Figure 7 and Table 3.

**Table 3.** Confusion matrix of classification using DBN, DDBN, 2parallel GDBN, LDA classifier.

| Method |  | Moving | Not Moving |
| --- | --- | --- | --- |
| DBN | Not Moving | 14% | 86% |
|  | Moving | 75% | 25% |
| DDBN | Not Moving | 23% | 77% |
|  | Moving | 93% | 7% |
| Two parallel GDBN | Not Moving | 15% | 85% |
|  | Moving | 97% | 3% |
| LDA classifier | Not Moving | 4% | 96% |
|  | Moving | 89% | 11% |

**Figure 6.**
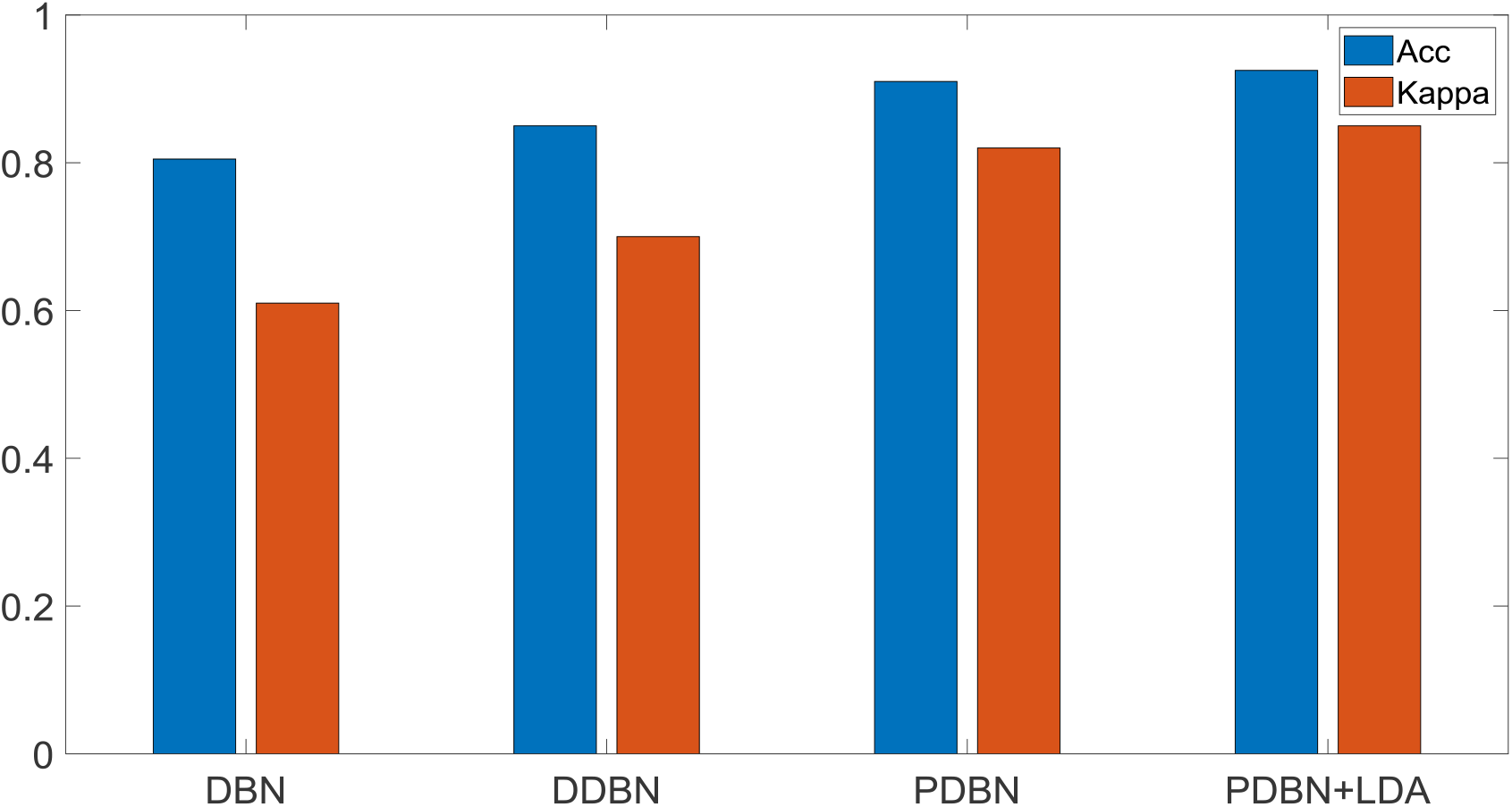
The test set accuracy and Kappa for the presented algorithms

**Figure 7.**
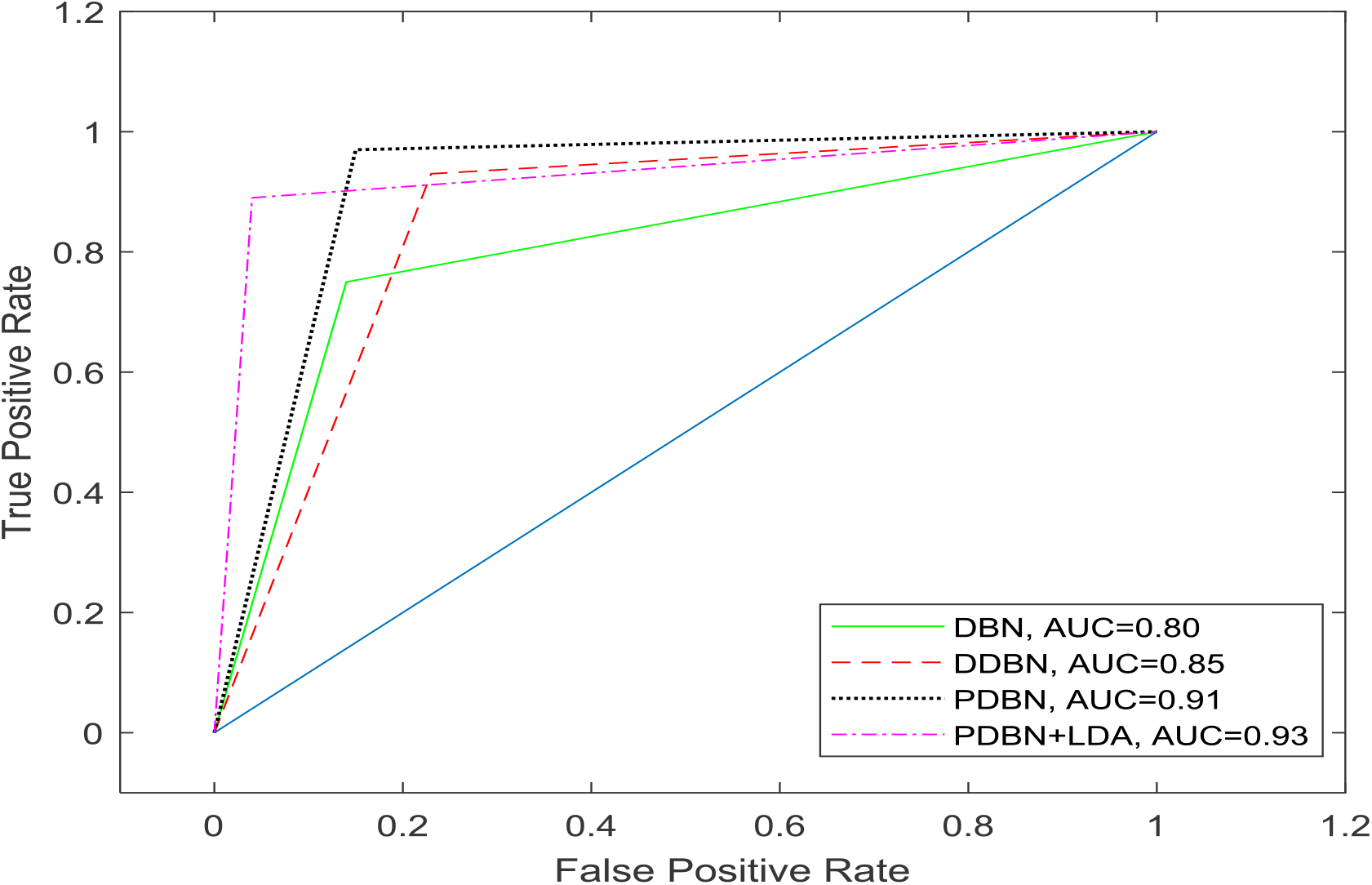
ROC curves for the classification results, using DBN method, DDBN, parallel Generative DBN, and LDA classifiers.

Figures 8 show the classification results in terms of real data and predicted data, using the different methods of RBM, DBN, Parallelizing Two Generative DBNs, and Cascading Two Parallelized GDBN to an LDA Classifier.

**Figure 8.**
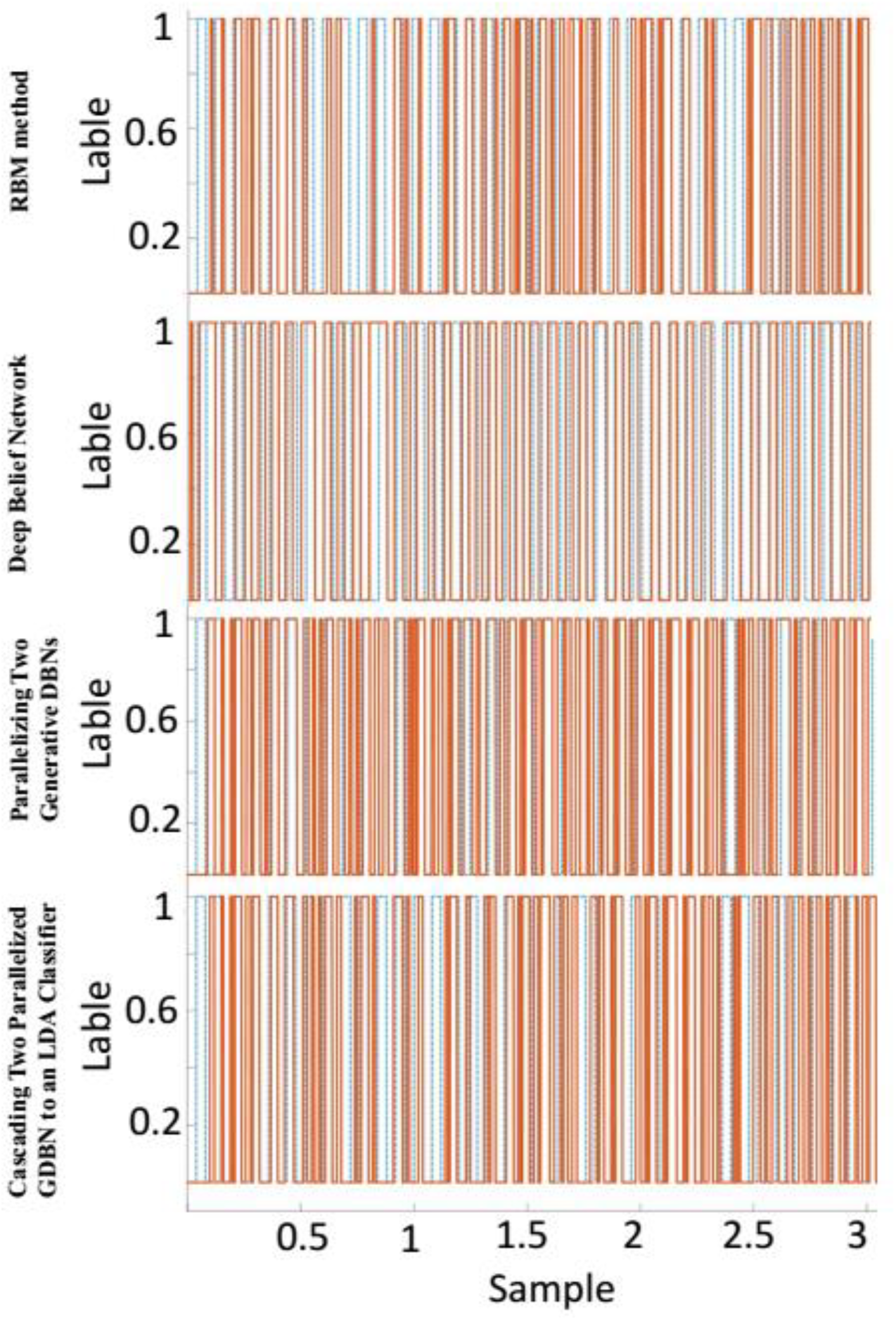
Classification results of Real and Predicted data, using different methods like RBM, DBN, Parallelizing Two Generative DBNs, and Cascading Two Parallelized GDBN to an LDA Classifier. The blue dash lines presented the real data and the red line showed the predicted data.

### Hybrid Classification

Using a combination of the EEG and EMG signals resulted in more accurate performance, especially if the disasters of patients are not related to their muscles such as aging, connate puniness of muscles, and fatigue. Initially, the intention recognized by the EMG signal. This process continued until the quality of the collected signals decreased. In this state, the supervisor control of the system switched to processing to EEG signal automatically. It was proofed that in the case of muscle fatigue drastic changes is some EMG signal characteristics suchlike Signal RMS, MPF, and FInsm are happened. Two discrete states can be considered for this system, the EMG based state and the EEG based state. The transition condition from a state to the other state is based on specific thresholds on the proposed parameters of EMG signal (Figure 9). The threshold of each index is a percentage of its initial value that presented in Table. The Simulink blocks based on RMS index are shown in Figure 9.

**Table 4.** Threshold of fatigue indices.

| Index | Threshold Percentage |
| --- | --- |
| RMS | 8% |
| MFP | 5% |
| FInsm | 12% |

**Figure 9.**
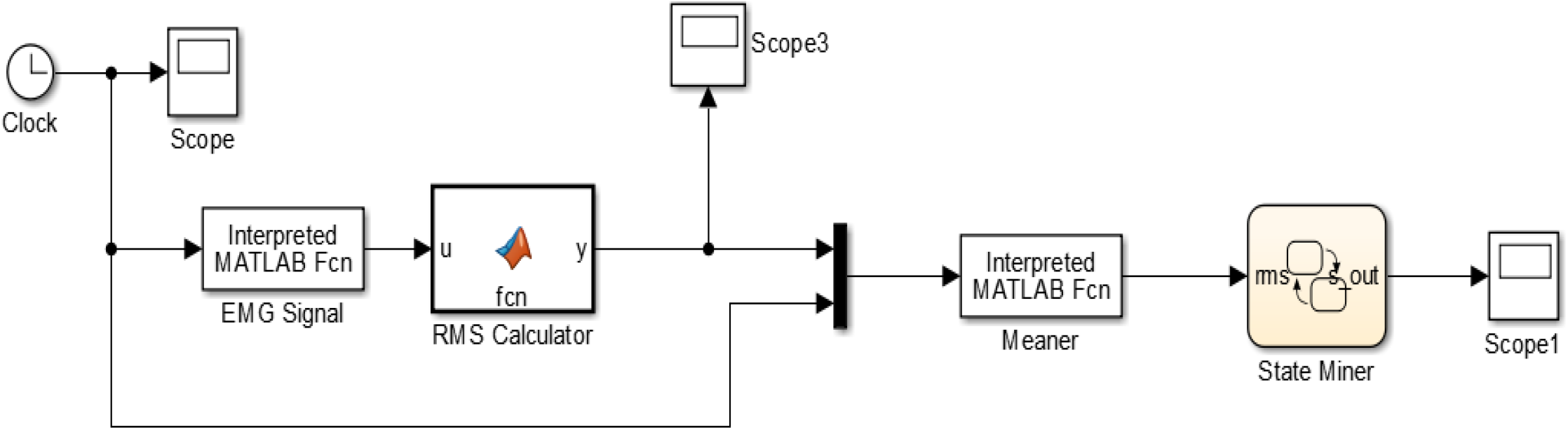
Block diagram of the EMG signal quality evaluator system implemented in SimuLink MATLAB

### Robustness of the Recognition System

It’s important to use a robust algorithm in order to recognize the intention of the user [Dehban, etal. 2015, 2016]. In order to test the robustness of the EEG processing methods, recorded signals from Ataxia patients were used (Figure 10). The most accurate algorithm in this case was parallel GDBNs using hard bound threshold which gives a 69% precision. It should be noted that the evaluation of this section had been done due to the number of movements in 400s.

**Figure 10.**
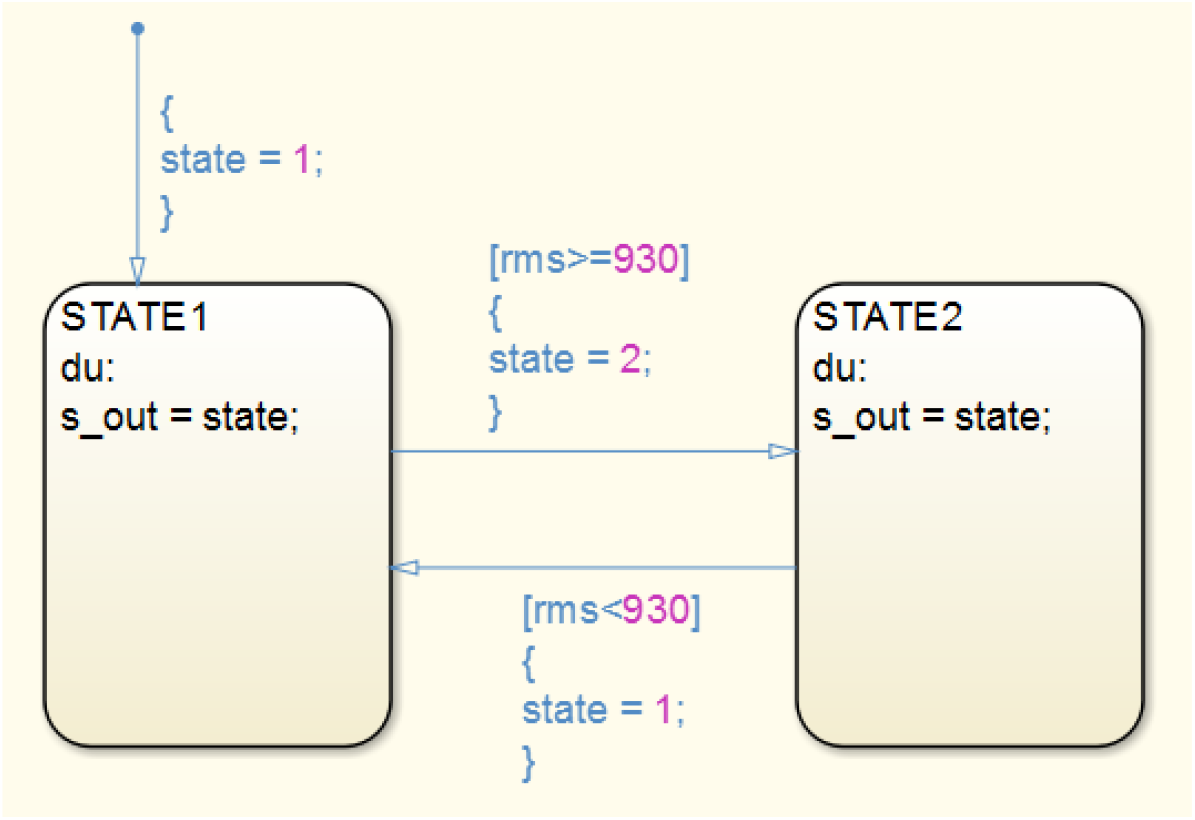
Hybrid dynamic system for state selection implemented through Stateflow(Matlab).

### Robotic Hardware Interface

A cable-driven robotic glove was fabricated as the hardware interface (Figure 10). It consists of five DC motors to pull the cables and create flexion movement. There are also five tension springs. Placed below the motors and connected to finger tips, by which fingers move back to their relaxed positions once the related motor is off.

DC motors are placed on back of the hand (dorsal aspect) using a rigid plate that is sewed to the glove. Back of the hand occurs minimum contacts during daily activities. Each motor is responsible for pulling and releasing the wires connected to its related finger. The Wires pass through adjusting pulleys that could be fixed along a rail. This allows us to adjust the tension in the wire, initial position of the fingers, range of finger motion or even to disable each motor individually. Each motor has two pulleys and two adjusting pulleys, one pulling the wire passed through all phalanges, which is responsible for flexion/extension movement, and the other one pulls the wire coming from proximal phalanx, which is responsible for adduction/abduction movement. Wires passes through rigid nodes, sewed to the glove at the middle of each phalanx, and then a plate on the palm of the hand that distributes the wires to their motors.

The wires move in a metal spring shield that protects them, and also allows them to move freely while holding an object. To perform flexion/extension, the adduction/abduction wires must be deactivated by adjusting their pulley position in the rail to lose the wires, and also adjust the pulleys of flexion/extension wires to make them tight. Multiple configurations and range of motion are achievable by just adjusting the pulleys position in their rail (Figure 11.A, 10.B).

**Figure 11.**
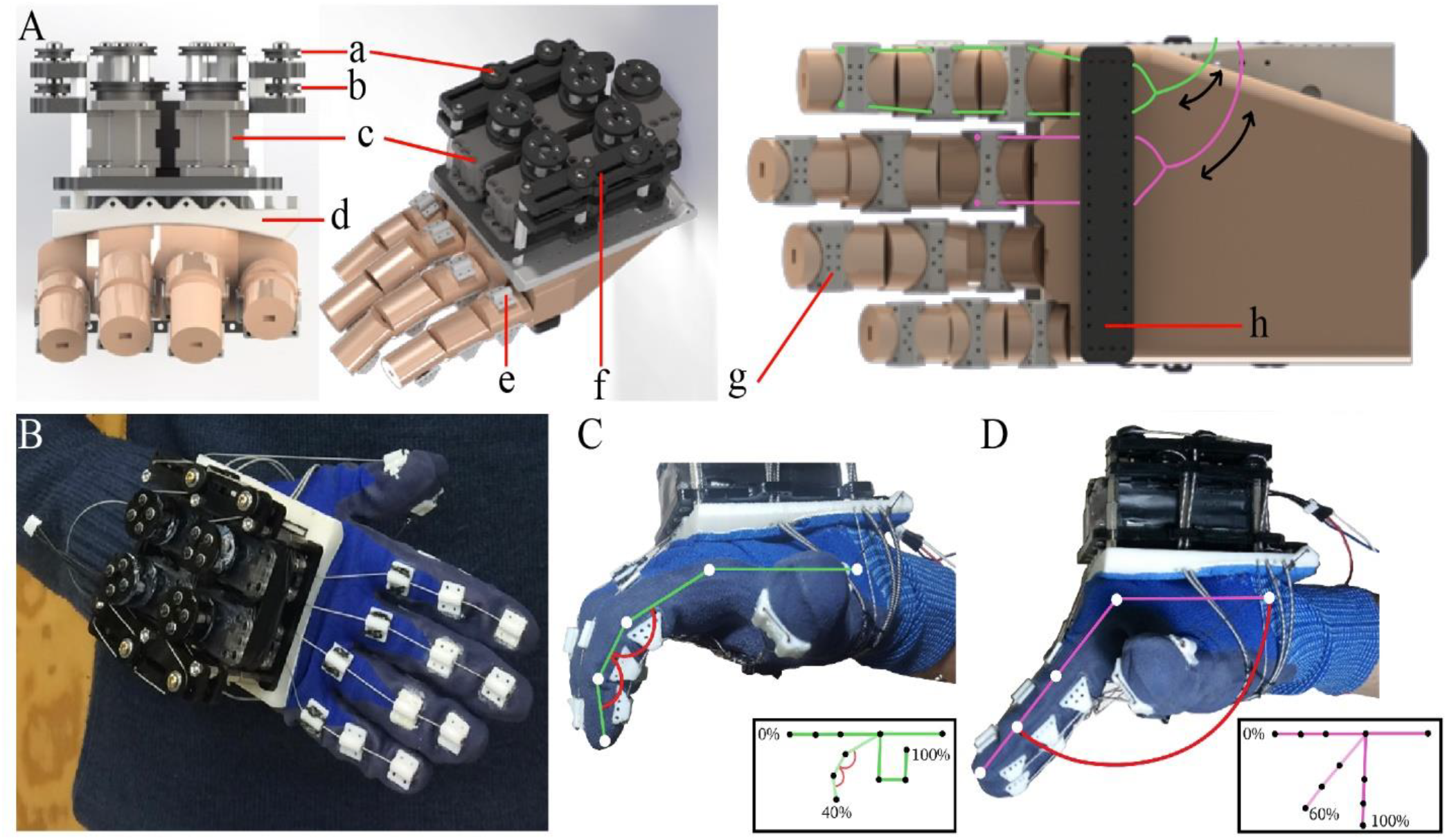
A) 3D CAD model of the robot. a) Adduction/abduction adjusting pulleys. b) Flexion/extension adjusting pulleys. c) Motors. d) rigid plat on back of the hand that holds all parts. e) Nodes on back of phalanges that guide the wires connected to springs for extension movement. f) adjusting rail. g) Nodes to guide flexion/extension and adduction/abduction wires. The wires for flexion movement shown in green pass all nodes on one finger and go to the motors by passing through their adjusting pulley (b), and the wires for abduction movement shown in purple pass only the proximal phalanx and go to the motors by passing through their adjusting pulley (a). h) Wire distributer placed on the palm. B) The first prototype of proposed robotic glove. C) Capability of the proposed robotic glove in flexion around DIP and PIP joints. D) Capability of the proposed robotic glove in flexion around MCP joints.

Initial results, achieved using image processing of the robot movements, shows that it can perform up to 40% of total flexion movement of the fingers around all three joints (Figure11.C) and up to 60% of the flexion movement around MCP joint (Figure11.D). It has been also demonstrated that rotation around DIP and PIP using the developed interface resembles the movement of a healthy finger (Figure 12). The robot includes some important features, such as low weight (maximum 2 kg), maximum degrees of freedom, and the ability to apply enough vertical force, portability and comfort.

**Figure12.**
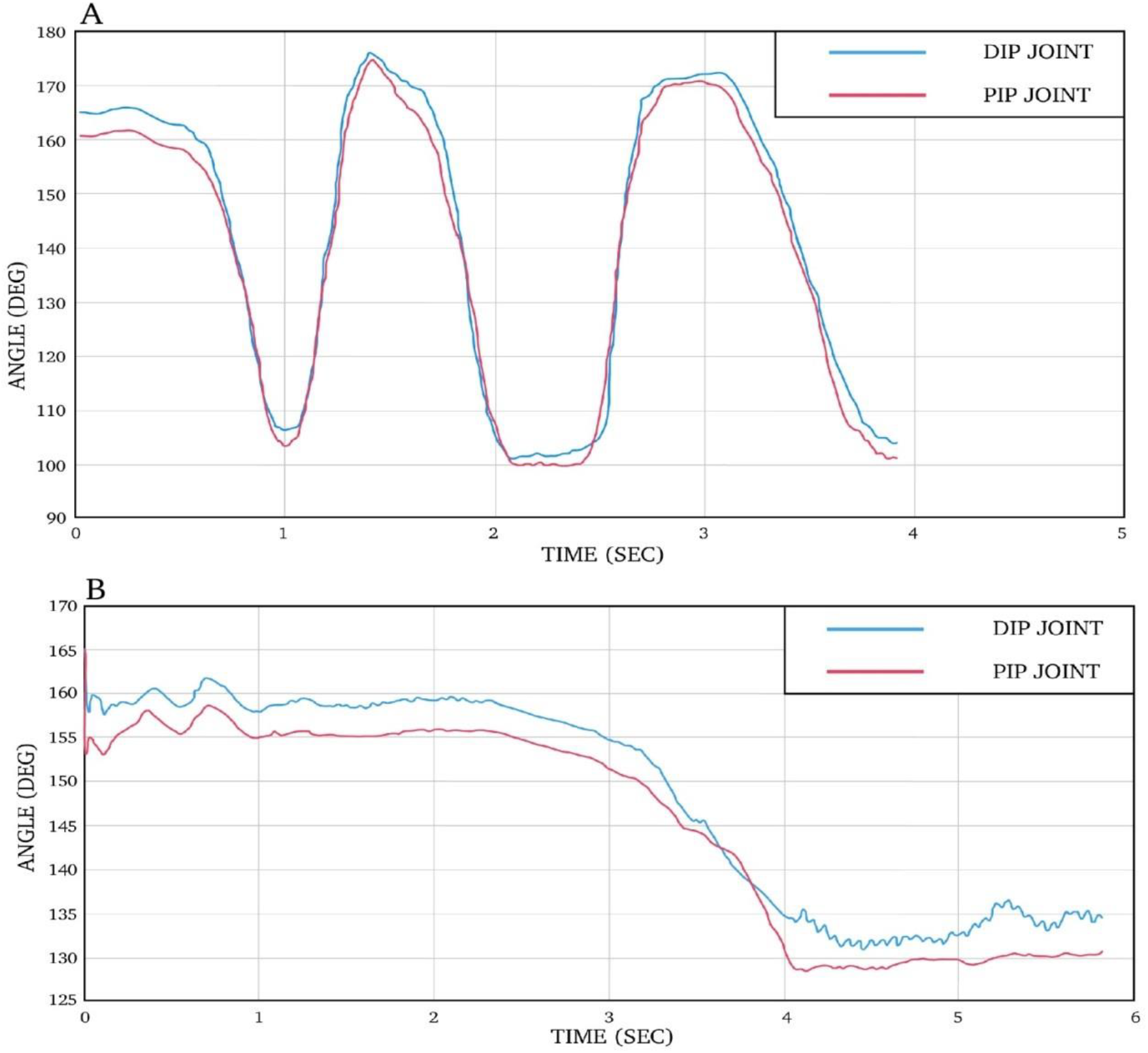
A) DIP and PIP angles of a real healthy hand in a normal flexion-extension movement B) DIP and PIP angles achieved by proposed robotic glove.

In addition, due to the low complexity of the robot, it is easy to work with and patients will not lose the desire to continue treatment. Our goal is to simplify the signals needed to actuate the robot and therefore move the fingers. In this case, each finger is connected to a DC motor that can be controlled by translated signals from our proposed EEG/EMG signal processing system.

## Discussion

In this study, a neuro rehabilitative hand exoskeleton was designed and developed for patients who have difficulty moving their fingers using a deep belief network mechanism. The wearable device has basic features such as small size and fit of the hand, comfortability, portability, ability to apply the appropriate force, understanding the decision of the patients to move the hand without the need of supervision in ADL.

In addition, we proposed a method to control the robot according to the user’s decision, through a human-machine interface. In this order, we compared different methods, and found that the serial auto encoder network with linear discriminant classification method and discriminative deep belief network had more accurate and acceptable results. Further with the aim of eliminating the training part before using the robot, the data of all healthy people were proceed using several deep belief networks.

Our results showed that the method of combining the two auto encoder networks reached the best accuracy. Then, the trained network was evaluated with the input of recorded data from the patients. It was observed that the decision to move in patients was recognized with a high accuracy. However, as expected, the results were not acceptable for patients with brain lesions. To solve this problem, a switching system was designed that also used muscle signals. Processing and detecting the decision to move using muscle signals is usually easier and reach a higher accuracy than the brain signal due to the distance from the sources of disturbance, the ability to track and be close to the target organ.

Therefore, in the proposed switching system, as long as the muscle signals are properly received, the control of the robot is lead through them, and as soon as its quality decreases, the system leads by the brain signals. The use of this assistive hybrid system, which has a higher recognition accuracy, is effective for patients whose motion problems are due to aging, small muscles or the like, in addition to patients suffering from spinal cord, cerebellum or nervous system lesions.

Furthermore, in order to develop a robot that is comfortable and light, this device was attached to a glove. The robot performs two flexion motions actively and the extension motions passively. In order to made the robot lighter, cable robots have been used instead of using rigid parts. Most cable robots have problems, such as thick cables, which limit the degree of freedom of the hand, and require powerful actuators as well as increasing the weight of the robot. In addition, due to the passing of cables from the back of the hand, there is no precise control on the motion of the fingers.

Many researchers developed 5-DOF EMG based wearable exoskeleton for post-stroke hand physical therapy [Hu, etal. 2013]. However, the device is bulky due to the rigid links and joints of the device which require accurate alignment with finger joints. In addition, another studies proposed a RoboGlove system and a haptic glove powered by a hydraulic system and artificial muscles. But, these devices were comparatively heavy [Hu, et al. 2013, Diftler et al. 2014]. In references [Borboni et al. 2018] and [Nycz, et al. 2016] researchers have developed an EMG based cable driven glove for hand rehabilitation. Another study developed a one DOF exoskeleton which is only applied to the index finger with an HMI that used brain and muscle signals to detect the user’s decision for moving the index finger [Guo, etal. 2016]. Some disadvantages of this system were the inability to be portable and also its degree of freedom. In [L. Randazzo, et al. 2017], researchers developed a BCI based portable wearable robot which is able to detect user’s decision to move each of the four fingers. Due to the use of a linear motor and its large size, which is located on the chest or back of the waist, power transmission cables are very large and as a result, it will be annoying for the patient.

In this study, by using thin cables and also changing the force point application of the cable on the palm, plus to weight loss, a suitable space was created for the free activity of the fingers and also did not restrict other degrees of hand freedom. To have a portable robot, dynamixel servomotors with high power, low dimensions and weight were selected. The processor was also chosen to be light in addition to real-time processing. The required power of the motors as well as the CPU must be supplied by the battery due to the portability of the robot. Other considerations such as the safety of the battery under pressure in certain conditions, lightness and no deformation over time are also considered.

One of the limitations of this project is that in order to be portable, light and compact, it was not possible to use powerful motors, and as a result, the system is not able to apply the maximum force of a healthy hand. Also, the cable mechanism is not well controlled because it has a lot of flexibility and therefore the design should be such that the device is in control as much as possible. In addition, although gloves are a suitable option for matching due to their high flexibility and elasticity, they lose their shape over time and the system performance is decreased. In the future, it is suggested to increase the degrees of freedom of the system for performing adduction and abduction motions. Also, to have a sense of touch, be equipped with appropriate sensors.

This paper presents an approach to take advantage of EMG/EEG-based approached signals to create a hybrid BCI device for controlling a hand exoskeleton through classifying the flexion attempt and resting states of the subjects. We analyzed data of 51 healthy and patients with brain lesions that involved the motor cortex and different hand movement disorders. The data are analyzed through different methods like deep neural network, restricted Boltzmann machine and LDA classifier methods to access a reliable accuracy. Then in order to evaluate the proposed approach, we designed and implemented a robotic system for rehabilitation of the hand movement to show that it is able to derive assistive control, by detecting flexion movement from the signals as a command and send it to robot. The accuracies with the combination of the EEG-EMG signals of healthies and patients were provided for comparison of effectiveness.

## Ethical Approval

Not required

## References

J. R. Wolpaw, N. Birbaumer, D. J. McFarland, G. Pfurtscheller, and T. M. Vaughan, “Brain–computer interfaces for communication and control,” Clinical neurophysiology, vol. 113, no. 6, pp. 767–791, 2002.

M. S. Al-Quraishi, I. Elamvazuthi, S. A. Daud, S. Parasuraman, and A. Borboni, “EEG-based control for upper and lower limb exoskeletons and prostheses: A systematic review,” Sensors, vol. 18, no. 10, p. 3342, 2018.

H. Gao et al., “EEG-based volitional control of prosthetic legs for walking in different terrains,” IEEE Transactions on Automation Science and Engineering, vol. 18, no. 2, pp. 530–540, 2019.

G. Kucukyildiz, H. Ocak, S. Karakaya, and O. Sayli, “Design and implementation of a multi sensor based brain computer interface for a robotic wheelchair,” Journal of Intelligent & Robotic Systems, vol. 87, no. 2, pp. 247–263, 2017.

K. Lee, D. Liu, L. Perroud, R. Chavarriaga, and J. d. R. Millán, “A brain-controlled exoskeleton with cascaded event-related desynchronization classifiers,” Robotics and Autonomous Systems, vol. 90, pp. 15–23, 2017.

Z. Li, J. Li, S. Zhao, Y. Yuan, Y. Kang, and C. P. Chen, “Adaptive neural control of a kinematically redundant exoskeleton robot using brain–machine interfaces,” IEEE transactions on neural networks and learning systems, vol. 30, no. 12, pp. 3558–3571, 2018.

D. P. Murphy et al., “Electroencephalogram-based brain–computer interface and lower-limb prosthesis control: A case study,” Frontiers in neurology, vol. 8, p. 696, 2017.

A. Prakash and S. Sharma, “A low-cost transradial prosthesis controlled by the intention of muscular contraction,” Physical and Engineering Sciences in Medicine, vol. 44, no. 1, pp. 229–241, 2021.

J. R. Wolpaw, D. J. McFarland, and T. M. Vaughan, “Brain-computer interface research at the Wadsworth Center,” IEEE Transactions on Rehabilitation Engineering, vol. 8, no. 2, pp. 222–226, 2000.

A. Chowdhury, H. Raza, Y. K. Meena, A. Dutta, and G. Prasad, “Online covariate shift detection-based adaptive brain–computer interface to trigger hand exoskeleton feedback for neuro-rehabilitation,” IEEE Transactions on Cognitive and Developmental Systems, vol. 10, no. 4, pp. 1070–1080, 2017.

R. Leeb, H. Sagha, R. Chavarriaga, and J. del R Millán, “A hybrid brain–computer interface based on the fusion of electroencephalographic and electromyographic activities,” Journal of neural engineering, vol. 8, no. 2, p. 025011, 2011.

R. Leeb, H. Sagha, and R. Chavarriaga, “Multimodal fusion of muscle and brain signals for a hybrid-BCI,” in 2010 Annual International Conference of the IEEE Engineering in Medicine and Biology, 2010: IEEE, pp. 4343–4346.

K. Kiguchi and Y. Hayashi, “A study of EMG and EEG during perception-assist with an upper-limb power-assist robot,” in 2012 IEEE International Conference on Robotics and Automation, 2012: IEEE, pp. 2711–2716.

A. Borboni et al., “Robot-assisted rehabilitation of hand paralysis after stroke reduces wrist edema and pain: a prospective clinical trial,” Journal of manipulative and physiological therapeutics, vol. 40, no. 1, pp. 21–30, 2017.

H. Mirzabagheiran, MB. Menhaj, AA. Suratgar and A. Sajedin, Temporal-Spatial Convolutional Residual Network for Decoding Attempted Movement Related EEG Signals of Subjects with Spinal Cord Injury, Computers in Biology and Medicine, 107159 2023.

L. Dovat et al., “A technique to train finger coordination and independence after stroke,” Disability and rehabilitation: Assistive technology, vol. 5, no. 4, pp. 279–287, 2010.

X. Huang, F. Naghdy, G. Naghdy, H. Du, and C. Todd, “The combined effects of adaptive control and virtual reality on robot-assisted fine hand motion rehabilitation in chronic stroke patients: a case study,” Journal of Stroke and Cerebrovascular Diseases, vol. 27, no. 1, pp. 221–228, 2018.

J. Iqbal, H. Khan, N. G. Tsagarakis, and D. G. Caldwell, “A novel exoskeleton robotic system for hand rehabilitation–conceptualization to prototyping,” Biocybernetics and biomedical engineering, vol. 34, no. 2, pp. 79–89, 2014.

Y. Fu, P. Wang, and S. Wang, “Development of a multi-DOF exoskeleton based machine for injured fingers,” in 2008 IEEE/RSJ International Conference on Intelligent Robots and Systems, 2008: IEEE, pp. 1946–1951.

A. Noormohamadi, MB. Menhaj, A. Sajedin, Control of leader–follower formation and path planning of mobilerobots using Asexual Reproduction Optimization (ARO), Applied Soft Computing, Elsevier. Vol. 14, 563–576, 2014.

M. A. Rahman and A. Al-Jumaily, “Design and development of a hand exoskeleton for rehabilitation following stroke,” Procedia Engineering, vol. 41, pp. 1028–1034, 2012.

S. Ueki et al., “Development of a hand-assist robot with multi-degrees-of-freedom for rehabilitation therapy,” IEEE/ASME Transactions on mechatronics, vol. 17, no. 1, pp. 136–146, 2010.

F. Vanoglio, A. Luisa, F. Garofali, and C. Mora, “Evaluation of the effectiveness of Gloreha (Hand Rehabilitation Glove) on hemiplegic patients. Pilot study,” in XIII congress of Italian Society of Neurorehabilitation, pp. 18–20, 2013.

P. Maciejasz, J. Eschweiler, K. Gerlach-Hahn, A. Jansen-Troy, and S. Leonhardt, “A survey on robotic devices for upper limb rehabilitation,” Journal of neuroengineering and rehabilitation, vol. 11, no. 1, pp. 1–29, 2014.

A. M. Khomami, F. Najafi, A survey on soft lower limb cable-driven wearable robots without rigid links and joints,“Robotics and Autonomous Systems”, vol.144, 2021

X. Hu, K. Tong, X. Wei, W. Rong, E. Susanto, and S. Ho, “The effects of post-stroke upper-limb training with an electromyography (EMG)-driven hand robot,” Journal of Electromyography and Kinesiology, vol. 23, no. 5, pp. 1065–1074, 2013.

M. Diftler et al., “RoboGlove-A robonaut derived multipurpose assistive device,” in International Conference on Robotics and Automation (ICRA), 2014.

C. J. Nycz, T. Bützer, O. Lambercy, J. Arata, G. S. Fischer, and R. Gassert, “Design and characterization of a lightweight and fully portable remote actuation system for use with a hand exoskeleton,” IEEE Robotics and Automation Letters, vol. 1, no. 2, pp. 976–983, 2016.

P. Fries, T. Womelsdorf T, R. Oostenveld, R. Desimone “The Effects of Visual Stimulation and Selective Visual Attention on Rhythmic Neuronal Synchronization in Macaque Area V4” 28 pp. 4823–4835, 2008.

A Gail, H Brinksmeyer, R. Eckhorn “Perception-related modulations of local field potential power and coherence in primary visual cortex of awake monkey during binocular rivalry”, Cereb. Cortex, 14: 300–313, 2004.

B. Pesaran, JS. Pezaris, M Sahani, PP. Mitra, RA Andersen, “Temporal structure in neuronal activity during working memory in macaque parietal cortex” 2002

A. Sajedin, MB. Menhaj, A-H. Vahabie, S. Panzeri, H. Esteky, “Cholinergic Modulation Promotes Attentional Modulation in primary visual cortex-A modeling study”. Scientific Reports vol 9(1), pp. 1–18, 2019.

U Mitzdorf, “Current source-density method and application in cat cerebral cortex: investigation of evoked potentials and EEG phenomena” Physiol. Rev 65: 37–99, 1985.

A. Sajedin, S. Salehi, H. Esteky, “Information Content and Temporal Structure of Face Selective Local Field Potentials Frequency Bands in IT Cortex”, Cerebral Cortex, 2024.

S. Guo, W. Zhang, J. Guo, J. Gao, and Y. Hu, “Design and kinematic simulation of a novel exoskeleton rehabilitation hand robot,” in 2016 IEEE International Conference on Mechatronics and Automation, 2016: IEEE, pp. 1125–1130.

L. Randazzo, I. Iturrate, S. Perdikis, and J. d. R. Millán, “mano: A wearable hand exoskeleton for activities of daily living and neurorehabilitation,” IEEE Robotics and Automation Letters, vol. 3, no. 1, pp. 500–507, 2017.

L.L.H.et al., “Effects of 8-week sensory electrical stimulation combined with motor training on EEG-EMG coherence and motor function in individuals with stroke”, Scientific Reports vol 8 (1) 2018.

Z. Wang, et al, Incorporating EEG and EMG patterns to evaluate BCI-based long-term motor training, IEEE Transactions on Human-Machine Systems 52 vol (4), 2022.

K Kiguch; Hayashi, Y., “Task Estimation of Upper-Limb Using EEG and EMG Signals”, IEEE/ASME Int. Conf. on Advanced Intelligent Mechatronics (AIM), 548, 2014.

A Dehban, MB Menhaj, A Sajedin, Neuro-ACT cognitive architecture applications in modeling driver’s steering behavior in turns, AUT Journal of Modeling and Simulation vol 47 (2), pp. 21–29, 2015.

A. Dehban, A. Sajedin, MB. Menhaj, Neuro-ACT Cognitive Architecture Applications In Modeling Driver’s Steering Behavior In Turns, Amirkabir International Journal of Science & Research (Electrical & Control Engineering) (AIJ-EEE), Vol. 45, 20–25, 2016.

J. Salgado Patrón.; C Barrera., “Robotic Arm Controlled By a Hybryd Brain Computer Interface”, ARPN Journal of Engineering and Applied Sciences, 11, 2016.

S. Tortora, L Toni., C Chisari, S Micera, E Menegatti, and F Artoni “Hybrid human-machine interface for gait decoding through Bayesian fusion of EEG and EMG classifiers”. Front. Neurorob. 2020.

J. Tryon, E Friedman, and A Trejos, “Performance evaluation of EEG/EMG fusion methods for motion classification,” in IEEE International Conference on Rehabilitation Robotics, pp. 971–976, 2019.

J. Tryon, and A Trejos, “Classification of task weight during dynamic motion using EEG-EMG fusion”. IEEE Sens. J. 21, 5012–5021, 2020.

H. Wöhrle, M Tabie, S Kim, K. Kirchner, and E Kirchner, “A hybrid FPGA-based system for EEG- and EMG-based online movement prediction”. Sensors 17:1552, 2017.

T. Dulantha Lalitharatne, K Teramoto, Y Hayashi, and K Kiguchi, “Towards hybrid EEG-EMG-based control approaches to be used in biorobotics applications: current status, challenges and future directions”. PALADYN J. Behav. Rob. 4, 147–154, 2013.

